# Explicit reward stabilizes motor output by attenuating sensory prediction error driven learning

**DOI:** 10.64898/2026.01.12.699038

**Authors:** Gaurav Panthi, Pratik Mutha

## Abstract

Motor adaptation driven by sensory prediction errors (SPEs) is often regarded as an automatic, implicit process that operates independent of reward. However, in most past work, reward and performance outcomes (success / failure) have been intrinsically confounded, making it unclear whether explicitly delivered reward *per se* influences SPE-driven learning. Here, we used an error-clamp paradigm to dissociate reward from both error and task outcome, enabling us to directly test whether explicit reward modulates implicit adaptation. Participants performed reaching movements under clamped visual feedback that produced a constant SPE, while being instructed to ignore the cursor. In two experiments, reward was delivered when the unseen *hand* successfully intersected the reach target. In an eight-target task, reward did not alter the overall magnitude or time course of adaptation. However, trial-level analyses revealed that rewarded movements were followed by smaller trial-to-trial updates, reduced variability, and a cumulative suppression of adaptive adjustments. Consistent with this result, individuals who experienced reward more frequently exhibited less asymptotic learning. In our second experiment using a simplified, two-target task, these trial-level effects accumulated to produce robust reductions in both asymptotic adaptation and aftereffects in the rewarded groups. Across both experiments, longer streaks of rewarded trials predicted progressively weaker SPE-driven updating. Collectively, these findings demonstrate that implicit adaptation is not insulated from reward signals. Instead, explicit reward appears to attenuate sensitivity to SPEs, stabilizing motor output particularly when the task structure allows consistent action-reward associations. We conclude that motivational signals can gate the expression of error-driven motor adaptation.

**SIGNIFICANCE STATEMENT:** Implicit motor adaptation is often thought to be insulated from motivational signals such as reward. By dissociating explicit reward from task outcome as well as visual error using an error-clamp paradigm, we show that reward reliably attenuates trial-to-trial adaptive corrections and reduces their variability, despite identical visual errors on every trial. These effects accumulate across rewarded movements and under stable task conditions, lead to reduced overall adaptation. Our findings thus demonstrate that reward can influence the expression of sensory error-based learning, and that task structure plays a critical role in revealing interactions between reward and implicit adaptation.

## INTRODUCTION

Reward is a fundamental determinant of behavior and learning, influencing how actions are selected, reinforced, and retained over time. In the motor domain, the effects of reward have been extensively studied within reinforcement-based paradigms, in which discrete signals (such as points, monetary gain, or success cues) inform participants about desired movement outcomes (Galea et al., 2015; Izawa & Shadmehr, 2011; Manley et al., 2014; Nikooyan & Ahmed, 2015; Vassiliadis et al., 2021). Across studies, reward has been shown to accelerate skill acquisition (Anderson et al., 2020; Dayan et al., 2014; Sporn et al., 2022), bias movement selection toward successful actions (Shmuelof et al., 2012; Yin et al., 2023), and enhance retention of newly learned motor behaviors (Abe et al., 2011; Galea et al., 2015; Huang et al., 2011; Vassiliadis et al., 2021). Critically, reward in several of these contexts typically functions as an independent driver of learning, capable of shaping behavior even in the absence of explicit error feedback (Izawa & Shadmehr, 2011; Nikooyan & Ahmed, 2015; van der Kooij & Overvliet, 2016).

In contrast, motor adaptation driven by sensory prediction errors (SPEs), or the mismatch between predicted and actual sensory outcomes of a movement, represents a distinct, largely automatic learning process (Hinder et al., 2010; Mutha et al., 2011; Shadmehr et al., 2010; Shingane et al., 2025; Tseng et al., 2007). In standard visuomotor adaptation tasks, SPEs are created by altering the relationship between the direction of hand motion and its feedback presented on a screen by means of a cursor. Strictly SPE-based adaptation is believed to be largely implicit, and results in updates to internal models without requiring conscious strategy or explicit awareness of the perturbation (Mazzoni & Krakauer, 2006; Morehead et al., 2017; Oza et al., 2024; Tsay et al., 2022). Whereas reward-based learning operates through reinforcement of successful outcomes, SPE-driven adaptation depends on the continuous recalibration of motor commands in response to discrepancies between predicted and observed sensory feedback. Importantly, these two forms of learning often co-occur in naturalistic tasks, yet the degree to which reward can influence adaptation that is *solely* SPE-driven remains poorly understood.

Bridging these domains, a small but growing body of work has examined whether signals typically associated with reward or task success modulate implicit motor adaptation. For example, Kim et al. (2019) and Oza et al. (2024) manipulated target size to vary the likelihood of the cursor hitting the target while keeping the SPE constant using “error-clamps”, in which visual feedback is constrained to follow a fixed trajectory relative to the target, independent of the participant’s actual hand movement. Participants who experienced more misses showed larger adaptive shifts than those whose movements were more often successful. Similarly, Al-Fawakhiri et al. (2023) used the error-clamp paradigm to examine the effect of target overlap and implicit task success on adaptation. Across these studies, adaptation was greater in the absence of reward-like or success cues, suggesting that reward or perceived success may suppress SPE-driven adaptation. However, a critical limitation of these experiments is that reward was never delivered explicitly. Instead, task success (cursor hitting the target) was operationalized as a proxy for reward. Because task outcome and reward were conflated, the specific influence of reward signals on implicit adaptation could only be inferred indirectly. In other words, even under clamped feedback, evaluative signals remained yoked to task outcome, such that successful and unsuccessful movements differed in their reward-like consequences despite identical SPEs. Consequently, these studies cannot determine whether explicitly-delivered reward itself, independent of task success or error magnitude, can modulate implicit adaptation.

Crucially, the error-clamp paradigm provides a means to address this gap by enabling a dissociation between error and evaluative feedback. Because the error is invariant and independent of hand motion, task success or failure is no longer informative about performance. This decoupling creates an opportunity to manipulate reward independent of both error and outcome, something that has not been adequately exploited in prior work. In the current study, we leverage this feature of the error-clamp paradigm to decouple SPEs from explicit reward delivery. By maintaining invariant feedback and manipulating reward independent of task success, learning can be driven solely by SPEs while evaluative feedback can be varied in an orthogonal manner. We consider two competing hypotheses. First, that reward stabilizes the adapted state (potentially through mechanisms involving dopaminergic modulation of cerebellar plasticity). Second, that implicit adaptation is insensitive to reward and reflects a process governed solely by SPEs that is independent of motivational or evaluative signals.

## METHODS

### Subjects

A total of 72 young, healthy, right-handed individuals (age range: 18–26 years, 20 female) were recruited for this study. Handedness was assessed using the Edinburgh Handedness Inventory (Oldfield, 1971). Participants were included only if they reported no neurological or cognitive disorders and no upper-limb injury. All procedures were approved by the Institutional Ethics Committee of the Indian Institute of Technology Gandhinagar, and all participants provided written informed consent. Subjects were monetarily compensated for their participation.

### Apparatus

Participants sat comfortably in an adjustable chair positioned in front of a Kinarm end-point robotic manipulandum (Kinarm, Ontario, Canada while holding its handle which permitted only planar arm movements (Figure 1a). The task display appeared on a HDTV mounted horizontally above the manipulandum. A mirror positioned between the screen and the robot reflected the visual display while occluding direct vision of the arm as well as the handle. Visual feedback about hand position was provided by means of an on-screen cursor. The motion of the cursor could be aligned with that of the true location of the hand or could be manipulated relative to hand motion.

**Figure 1.**
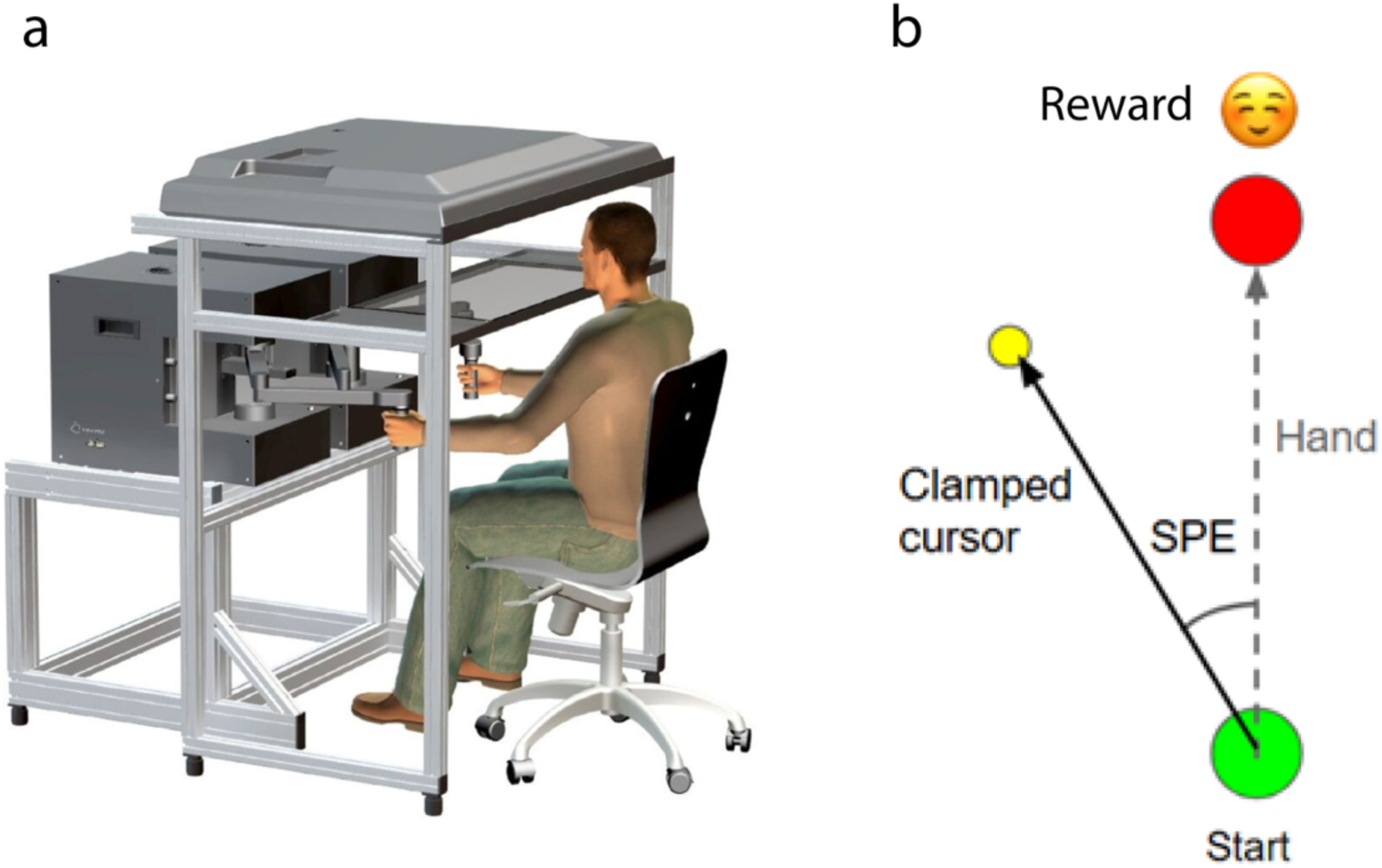
**(a)** Experimental Setup. Participants performed reaching movements while holding the handle of a robotic manipulandum (Kinarm end-point). The start position, targets, and cursor feedback were displayed on a screen and reflected onto a mirror placed horizontally between the screen and the hand, occluding direct vision of the arm. **(b)** Experimental Task. Participants made reaching movements from start position (green circle) toward the target (red circle). Movement feedback was provided by means of a cursor (yellow circle), but its motion was clamped at a 30° offset from the hand motion. Subjects were instructed to ignore the invariant cursor and move their hand through the target. If the hand successfully intersected the target, a smiley face emoji was displayed as a reward signal for the “visual reward” group. For the “visuoproprioceptive reward” group, the smiley was paired with a haptic event at the hand. A third, “control” group, received no reward.

### Task design

Participants made fast point-to-point movements from a circular start position (1 cm radius) to a target displayed on the screen. Each trial began when the cursor was brought into, and remained within, the start position for 500 ms; premature exit from the start position reset the trial. After the hold period, a circular target (1 cm radius) appeared at a distance of 12 cm, and subjects were instructed to make a rapid, slicing reach through the target. During the reach, one of three types of cursor feedback was provided: 1) veridical feedback wherein the cursor accurately reflected hand position, 2) no feedback wherein the cursor was invisible, or 3) error-clamped feedback where the cursor followed an invariant trajectory rotated by a fixed angle, independent of the actual hand movement. Across both our experiments, the clamped cursor was rotated by 30° counterclockwise from the line connecting the start position and the target. On the clamp trials, participants were clearly instructed and made to understand that they should ignore the cursor and aim their hand directly toward the displayed target.

### Experiment 1

In experiment 1, participants (n = 36) were randomly assigned to one of three groups: control, visual reward, or visuoproprioceptive reward (n = 12 each). All participants, regardless of the group they belonged to, performed 240 reaches to one of eight targets arranged radially from the start position at 0°, 45°, 90°, 135°, 180°, 225°, 270°, and 315°. Targets were presented pseudorandomly. The experiment comprised three consecutive blocks: baseline, learning, and washout. After a brief familiarization phase, participants completed a 40-trial baseline block consisting of two sub-blocks of 16 no-feedback trials followed by 24 veridical-feedback trials. They then completed 160 error-clamp learning trials in which the cursor followed a fixed 30° rotated trajectory. This produced a constant SPE, as participants aimed their hand toward the target but visual feedback deviated from it. Instructions to ignore the cursor and aim the hand toward the target were given prior to the start of this learning block. The washout block mirrored the baseline structure: 16 no-feedback trials followed by 24 veridical-feedback trials. Before the start of the washout block, subjects were again instructed to aim their hand directly toward the target.

The two reward groups had identical baseline and washout blocks but differed in terms of the reward received during learning. Notably, reward was delivered whenever the (invisible) *hand* intersected the target. In the visual reward group, a smiley face appeared on the screen for 1s just above the target (Figure 1b). In the visuoproprioceptive reward group, the smiley feedback was paired with a haptic event associated with the target. Specifically, we implemented a virtual force ring (0.1 cm thickness) around the target with 1000 N/m stiffness. When the participant’s hand entered the target region, a velocity-dependent damping force was applied, producing a cushion-like sensation that gave the impression of contacting a compliant physical object. This “dual” reward enabled us to assess whether additional reward-related information conveyed haptically would strengthen the reward effects. Note that the reward was given when the hand successfully entered the target region, while the SPE induced due to the cursor clamp would tend to pull the hand away from it. Thus, during the learning block, reinforcement-mediated behavior and SPE-based adaptation were essentially in conflict. The control group did not receive any reward.

### Experiment 2

In Experiment 2, we again recruited 36 participants, and assigned them to three groups (n = 12 each). The overall trial structure was similar to Experiment 1, except that only two targets (located at 90° and 270°) were presented. The baseline block again consisted of 16 no-feedback trials followed by 24 veridical-feedback trials. The learning block used the same 30° error clamp, producing a consistent SPE, and participants were instructed to ignore the cursor and move their hand directly toward the target. As before, reward was delivered in the visual and visuoproprioceptive reward groups when the hand successfully intersected the target. In the visual reward group, a smiley emoji was displayed for 1s; in the visuoproprioceptive group, the smiley was paired with the haptic event as described in Experiment 1. The control group received no reward. The reduced target set (only 2 targets here in Experiment 2 versus 8 in Experiment 1) allowed us to evaluate the influence of reward under a simpler task structure requiring fewer distinct movement plans.

### Data Analysis

Hand (robot endpoint) position data were sampled at 1000 Hz and filtered using a 10 Hz Butterworth filter. Velocity was obtained by differentiating the filtered position signal, and the point of peak velocity was identified on each trial. The primary dependent variable was the hand deviation, defined as the angle between the vector from the start position to the target and the vector from the start position to the hand position at peak velocity. To correct for baseline biases, the mean hand angle from all trials of the baseline phase was subtracted from the hand deviation on each trial. Trials on which no movement was initiated, or where hand deviation exceeded ±100°, were excluded. To further filter out outliers that fell within the ±100° range, we applied a moving average filter with an 8-trial window and a 2.5 standard deviation threshold. In total, less than 1% of trials were excluded across both experiments together.

To assess how learning evolved across trials, we defined “early” learning as the mean hand angle calculated over the first 32 learning trials. Likewise, “final” learning was defined as the mean hand deviation over the last 32 learning trials. Aftereffects were defined as the mean hand deviation during the first 16 no-feedback trials of washout. We also calculated single-trial learning (STL) as the trial-to-trial change in hand deviation. This was done by subtracting the hand deviation on the (n+1)^th^ trial from that on the n^th^ trial. The hand deviation and single trial learning were subjected to various statistical tests, presented with the corresponding results below. The significance threshold was set at 0.05.

## RESULTS

### Experiment 1

In the first experiment, three groups of participants performed the learning trials under clamped visual feedback conditions, with the instruction to ignore this feedback and move their hand directly to the target. For two of the three groups, an explicit reward signal was delivered when their hand successfully intersected the target (Figure 1b). One group received a visual reward in the form of a smiley emoji (visual reward group), while the other received the same visual feedback along with a soft force pulse to the hand (visuoproprioceptive reward group). The third group received no reward and served as a control.

Figure 2a shows the change in hand angle across cycles for all three groups (1 cycle = 8 trials). All subjects displayed the canonical pattern of error-clamp adaptation; the hand drifted gradually opposite to the direction of the clamped cursor, eventually reaching a stable asymptote. Consistent with prior work, this drift occurred despite instructions to ignore the cursor. No group differences were evident in the overall learning curves.

**Figure 2.**
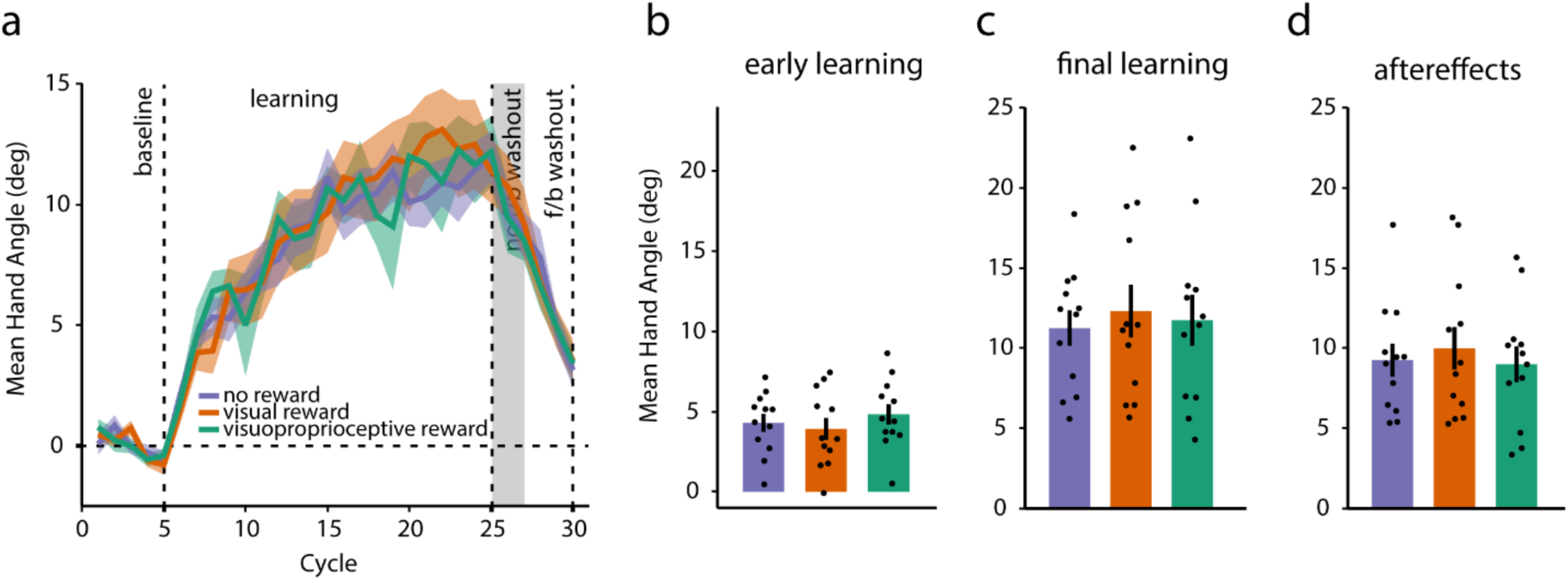
**(a)** Group-averaged hand deviations across cycles for the three experimental groups. Shaded regions represent SEM. **(b–d)** Mean ± SEM hand deviation during **(b)** early learning, **(c)** final learning, and **(d)** no-feedback washout phase. Dots represent individual participants.

During early learning (Figure 2b), the mean ± SEM hand-angle deviations were 4.297 ± 0.554° (control), 3.917 ± 0.687° (visual reward), and 4.832 ± 0.628° (visuoproprioceptive reward). These differences were not significant in a one-way ANOVA (F_2,33_ = 0.54, p = 0.588, 1^2^_p_ = 0.032). As learning progressed, all groups reached comparable asymptotic levels (Figure 2c; control = 11.253 ± 1.098°, visual reward = 12.315 ± 1.633°, visuoproprioceptive reward = 11.747 ± 1.589°), with no reliable group differences evident at final learning (F_2,33_ = 0.132, p = 0.876, 1^2^_p_ = 0.008). Aftereffects measured during the no-feedback washout trials (Figure 2d) were also statistically indistinguishable across groups (F_2,33_ = 0.201, p = 0.819, 1^2^_p_ = 0.012). Separate independent samples t-tests comparing the two rewarded groups showed no differences at either final learning (t_22_ = 0.25, p = 0.805) or during washout (t_22_ = 0.579, p = 0.568). Exponential fits of individual learning curves revealed no between-group differences in learning rate either (F_2,33_ = 0.795, p = 0.46, 1^2^_p_ = 0.046). Together, these analyses indicated that explicit reward did not affect the overall magnitude or the time course of implicit adaptation in this multi-target environment.

Although the group-level learning curves showed no reward effect, we probed whether reward might nevertheless modulate *trial-to-trial* learning (single trial learning, STL). To probe this, we modeled the magnitude of the update on trial *n+1* as a function of whether trial *n* was rewarded using a linear mixed-effects model with a fixed effect of reward and random intercepts for participants. The model was set up as follows:

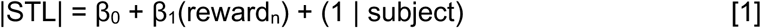

Here, β_0_ represents the update magnitude following unrewarded trials whereas β_1_ captures the change in update magnitude following rewarded trials. A negative β_1_ indicates reward-related suppression of the subsequent implicit adjustment. We found that in the visual reward group, reward significantly reduced STL magnitude (β_1_ = −1.19°, 95% CI = [−1.98°, −0.41°], p = 0.0028), with updates being approximately 12% smaller following rewarded trials. A similar pattern was observed in the visuoproprioceptive reward group (β_1_ = −1.82°, 95% CI = [−2.73°, −0.93°], p < 0.0001), where updates following rewarded trials were about 18% smaller than those following unrewarded trials. Thus, even though the visual error was identical on every trial due to the clamp, the presence of reward on trial *n* attenuated the magnitude of the update on trial *n+1*.

Given that rewarded trials produced smaller single-trial updates than unrewarded trials in both reward groups, we next asked whether this suppressive effect scaled with the accumulation of reward across consecutive movements. For each learning trial, we computed a reward “streak”, defined as the number of consecutively rewarded trials immediately preceding the current trial. Because long streaks occurred idiosyncratically, we restricted this analysis to streaks that appeared at least twice within a subject and were observed in at least three subjects. Under these criteria, the obtained streak length was 5 in the visual reward group and 4 in the visuoproprioceptive reward group (Figure 3a). This preserved the reliably sampled portion of the streak distribution while ensuring that each included streak level reflected reproducible behavior rather than any outliers.

**Figure 3.**
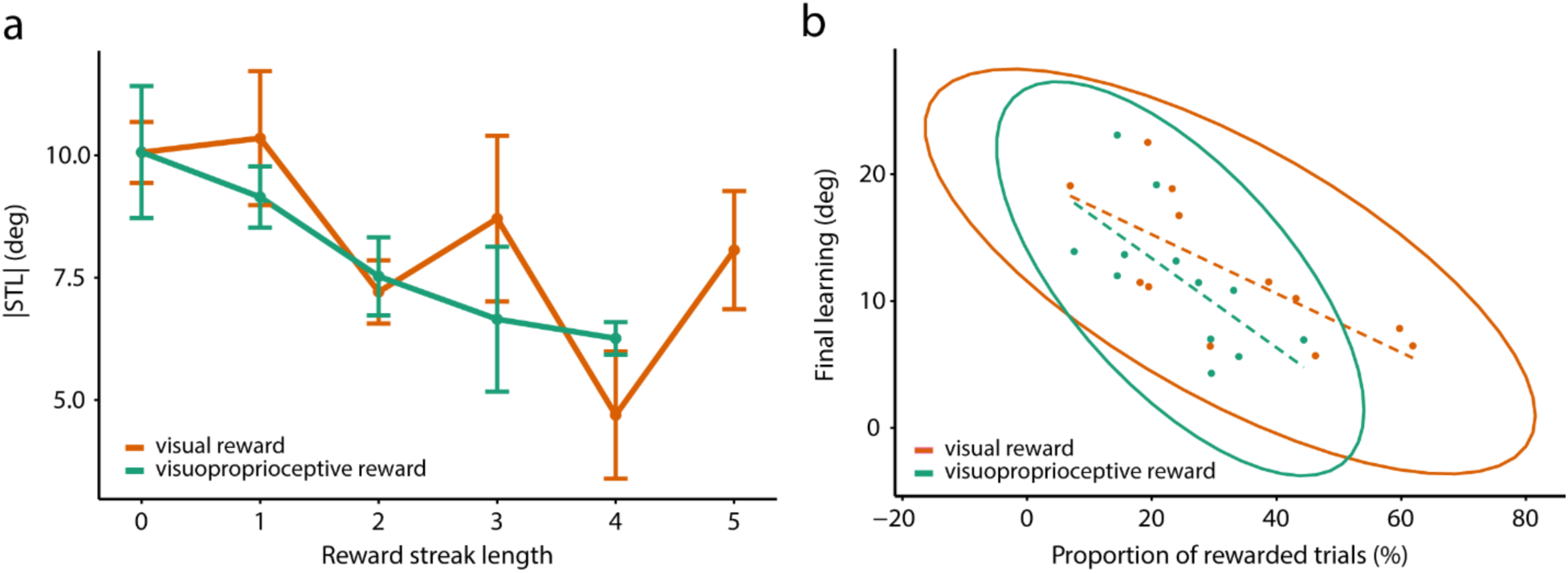
**(a)** Single trial learning as a function of reward streak length for Experiment 1. Error bars represent SEM. The magnitude of STL decreased as streak length increased. **(b)** Final learning level as a function of proportion of rewarded trials for the two reward groups. Ellipses are 95% confidence ellipses and the lines represent the linear regression fit to the data for each group. Dots are individual subjects.

We then examined whether streak length predicted the magnitude of the subsequent update using the following linear mixed-effects model, fit separately for the visual and visuoproprioceptive reward groups:

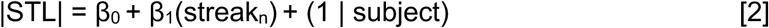

Here, β_1_ captures the change in SPE-driven updating associated with each additional rewarded trial in the streak. In both reward groups, β_1_ was significantly negative (visual reward: β_1_ = −0.64°, 95% CI = [−1.19°, −0.09°], p = 0.0259; visuoproprioceptive reward: β_1_ = −1.005°, 95% CI = [−1.54°, −0.467°], p = 0.0008, Figure 3a), indicating that longer sequences of rewarded movements progressively reduced the magnitude of the subsequent adaptive correction. Thus, reward did not merely attenuate single adaptive responses, but accumulated across trials to suppress the ongoing expression of SPE-driven implicit learning.

If reward transiently attenuates single-trial learning, then participants who experience reward more often should exhibit smaller overall adaptation. To test this, we correlated each participant’s proportion of rewarded trials with their final adaptation magnitude. Across participants, reward frequency correlated negatively with final learning (visual reward: *r* = −0.713, 95% CI = [−0.913, −0.236], p = 0.0092; visuoproprioceptive reward: *r* = −0.667, 95% CI = [−0.897, −0.151], p = 0.0178, Figure 3b). Thus, individuals who received reward more frequently tended to exhibit reduced asymptotic learning, consistent with the cumulative consequences of reward-driven suppression of single-trial corrections.

Finally, we assessed whether reward influenced the variability of trial-to-trial adjustments. For each participant, we computed the standard deviation of STL separately for trials that followed reward and for those that followed no-reward, and compared these two variance measures using paired t-tests within each reward group. In the visual reward group, variability after rewarded trials was significantly lower than after unrewarded trials (t_11_ = −7.85, p < 0.001); a similar pattern was observed in the visuoproprioceptive reward group as well (t_11_ = −4.004, p = 0.002). This indicated that rewarded movements were followed by more consistent, less variable adaptive corrections, complementing the reduction in update magnitude observed in the single-trial analyses.

In sum, Experiment 1 revealed that while explicit reward did not alter the overall magnitude or time course of implicit adaptation, it attenuated trial-to-trial learning. Rewarded trials, especially sequences of rewarded trials, produced smaller adaptive corrections and reduced variability, and participants who experienced reward more frequently ultimately exhibited less net adaptation. Nonetheless, given that reward exerted robust trial-level effects but did not modulate overall learning in this 8-target design, we asked whether reducing target variability might enhance the expression reward effects on SPE-driven adaptation. Experiment 2 therefore used a simplified 2-target design to test whether a more stable movement context would amplify reward-related differences in implicit learning.

### Experiment 2

In our second experiment, we asked whether reducing task complexity would reveal an influence of reward on implicit adaptation that was not detectable in the 8-target design. As in Experiment 1, participants reached under clamped visual feedback, but here movements were made to only two targets located 180° apart. This design provided more repetitions per target, thereby increasing the likelihood that reward could influence the stability of implicit adaptation and consistently reinforce the same motor plan.

Figure 4a shows the evolution of hand angles across learning cycles for the three groups (1 cycle = 8 trials). As in Experiment 1, all groups exhibited the characteristic drift opposite to the clamped direction, reflecting SPE-driven adaptation. During early learning (Figure 4b), hand-angle deviations were comparable across groups as revealed via a one-way ANOVA (control = 6.388 ± 1.045°, visual reward = 5.578 ± 0.813°, visuoproprioceptive reward = 6.573 ± 0.885°; F_2,33_ = 0.332, p = 0.72, 1^2^_p_ = 0.02). However, in contrast to Experiment 1, clear group differences emerged as learning progressed. By the end of learning (Figure 4c), the control group exhibited significantly larger asymptotic adaptation (12.548 ± 1.811°) than both reward groups (visual reward = 7.733 ± 0.945°, visuoproprioceptive reward = 7.294 ± 1.135°; F_2,33_ = 4.67, p = 0.016, 1^2^_p_ = 0.221). Post-hoc analyses using Dunnett’s test confirmed that both reward groups differed significantly from control (visual reward: p = 0.031 and visuoproprioceptive reward: p = 0.018). A similar pattern was evident during washout (Figure 4d), where aftereffects were significantly different across groups (F_2,33_ = 5.206, p = 0.011, 1^2^_p_ = 0.240). Again, Dunnet’s post-hoc test indicated that the control group had larger aftereffects than both reward groups (visual reward: p = 0.04 and visuoproprioceptive reward: p = 0.008). These findings indicated that explicit reward attenuated the overall expression of SPE-driven adaptation when the task involved only two targets.

**Figure 4.**
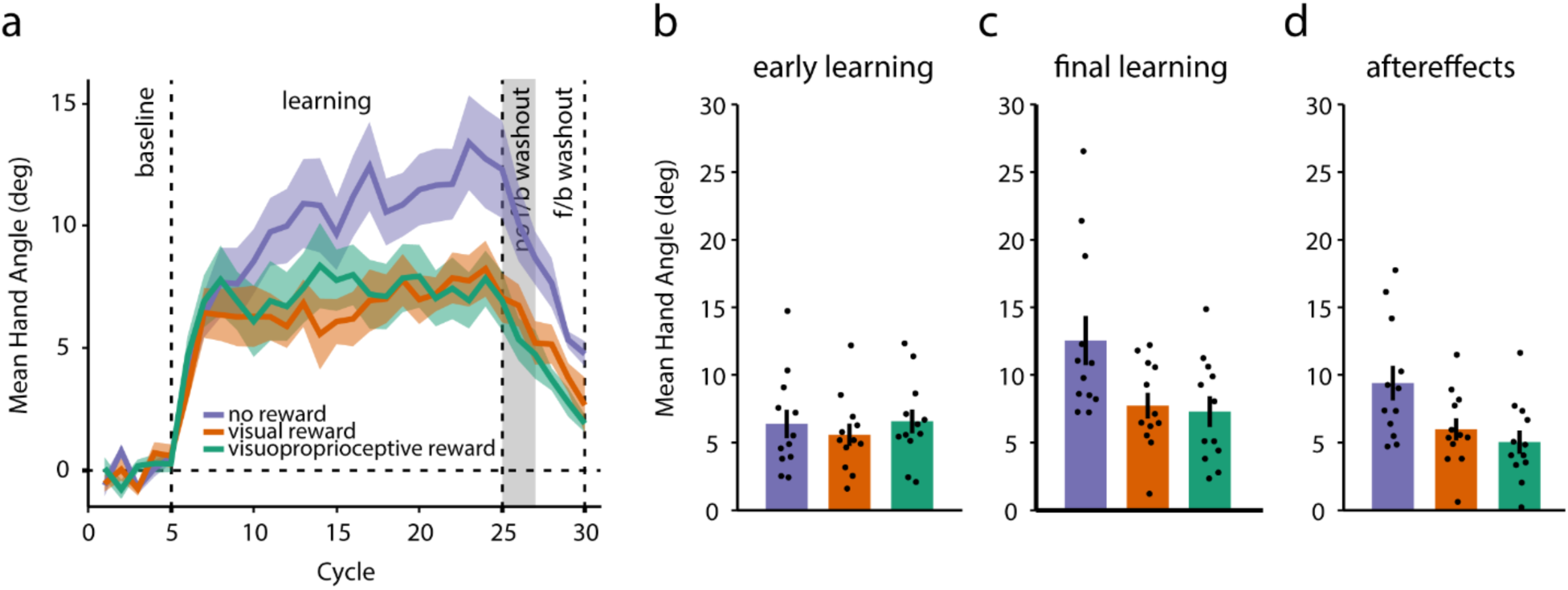
**(a)** Group-averaged hand deviations across cycles for the three experimental groups. Shaded regions represent SEM. **(b–d)** Mean ± SEM hand deviation during **(b)** early learning, **(c)** final learning, and **(d)** no-feedback washout phase. Dots represent individual participants.

To examine whether reward influenced the trial-to-trial updating of motor output, we again analyzed single-trial learning (STL), defined as the change in hand angle from trial *n* to *n+1*. We applied the linear mixed-effects model (equation 1) used in Experiment 1 and observed that consistent with the block-level findings, β_1_ was significantly negative for the visual reward group (β_1_ = −1.53°, 95% CI = [−2.12°, −0.94°], p < 0.0001) as well as the visuoproprioceptive reward group (β_1_ = −0.66°, 95% CI = [−1.202°, −0.118°], p = 0.017). This indicated that reward tended to attenuate SPE-driven updating on a trial-to-trial basis.

We next examined how reward accumulation across consecutive trials (streaks) influenced single-trial learning. As before, to ensure that each included streak level was supported by stable, non-idiosyncratic data, for each participant, we included only those reward streak lengths that occurred at least twice within a participant and were represented in at least three participants within the group. This yielded a streak length of 13 in the visual reward group and 12 in the visuoproprioceptive reward group (Figure 5a). Using these streak values as predictors of trial-to-trial update magnitude (equation 2), we found a consistent negative relationship between reward accumulation and STL. In the visual reward group, longer streaks were associated with significantly smaller updates (β_1_ = −0.129°, 95% CI = [−0.227°, −0.03°], p = 0.012); a similar effect emerged in the other reward group (β_1_ = −0.134°, 95% CI = [−0.235°, −0.033°], p = 0.011). Thus, a richer reward history reliably predicted weaker SPE-driven updating.

**Figure 5.**
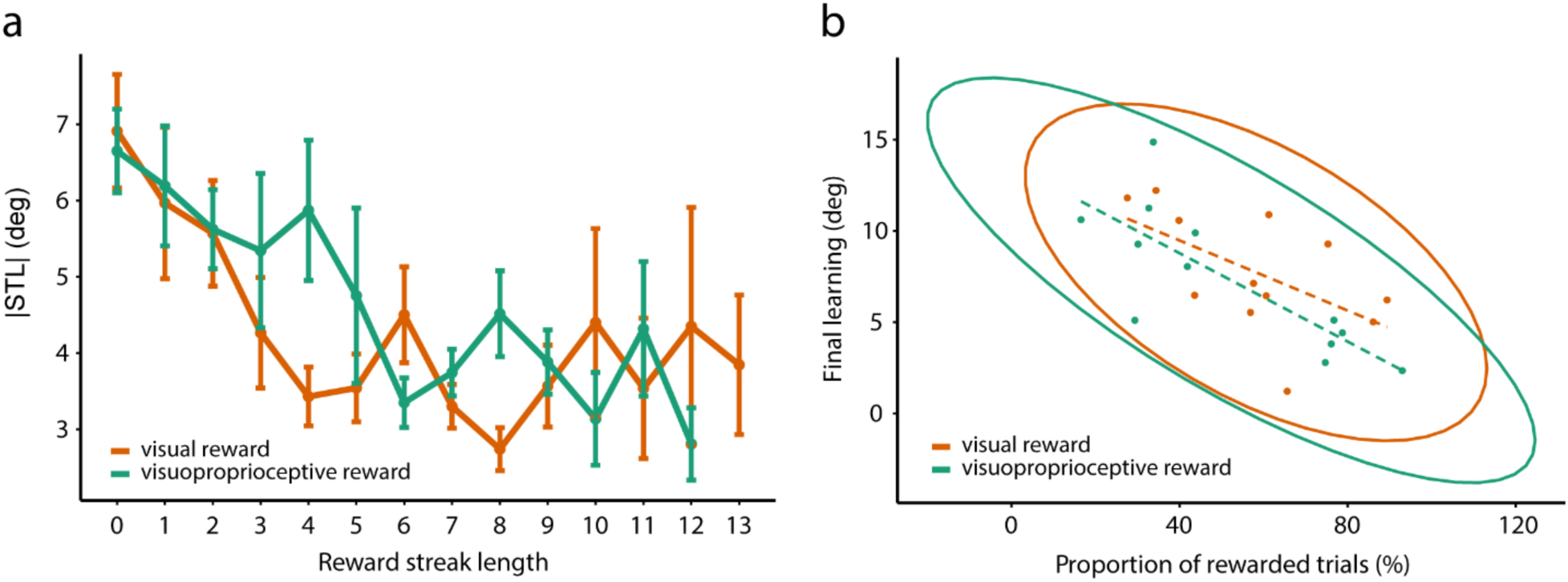
**(a)** Single trial learning as a function of reward streak length for Experiment 2. For both groups, STL magnitude decreased as streak length increased. Error bars represent SEM. **(b)** Final learning level as a function of proportion of rewarded trials for the two reward groups. Ellipses are 95% confidence ellipses, and the lines represent the linear regression fit to the data for each group. Dots are individual subjects.

Given this trial-level attenuation, we next tested whether the extent of reward exposure predicted each participant’s final level of adaptation. For both reward groups, the proportion of trials on which participants received reward was negatively correlated with their final learning magnitude (visual reward: r = −0.571, 95% CI = [−0.863, 0.004], p = 0.05; visuoproprioceptive reward: r = −0.789, 95% CI = [−0.938, −0.394], p = 0.002; Figure 5b). Thus, participants who were rewarded more frequently exhibited smaller asymptotic adaptation, mirroring the effect observed in Experiment 1, but more aligned with group-level differences.

Lastly, we examined whether reward altered the variability of trial-to-trial adjustments. For each participant, we computed the standard deviation of STL separately for rewarded and unrewarded trials and compared these values using paired t-tests. In the visual reward group, variability was significantly lower following rewarded trials (t_11_ = −2.785, p = 0.018), indicating that reward stabilized motor output by reducing trial-to-trial fluctuations. In the visuoproprioceptive reward group, while variability following rewarded trials was also smaller compared to the unrewarded trials, the difference did not reach statistical significance (t_11_ = −1.859, p = 0.09).

Taken together, the findings from Experiment 2 brought forth more clearly the influence of reward on implicit adaptation. In this simplified context, reward reduced the influence on sensory prediction errors, weakening both trial-by-trial adaptive adjustments and the ultimate extent of adaptation. These converging effects suggest that reward interacts with implicit learning when movements are repeated in a sufficiently stable manner, revealing a suppressive influence that was masked in the more variable, multi-target environment of Experiment 1.

## DISCUSSION

The present study examined whether explicitly delivered reward modulates implicit sensorimotor adaptation. Implicit adaptation has traditionally been considered to be an automatic process driven solely by SPEs (Kumar et al., 2020; Morehead et al., 2017; Oza et al., 2024; Tsay et al., 2022; Tseng et al., 2007). While recent work has suggested that reward may interact with error-based learning (Forano & Franklin, 2024; Izawa & Shadmehr, 2011; Nikooyan & Ahmed, 2015; Roth et al., 2024; van der Kooij & Overvliet, 2016), many of these studies have conflated reward with task outcome (successfully hitting a reach target), making it difficult to determine whether reward *per se* influences implicit learning. To address this issue, here we used clamped error feedback to induce learning while also providing explicit reward that was independent of the task outcome. Across two experiments, we found that explicit reward signals systematically attenuated trial-to-trial adaptation and reduced the overall expression of learning. This attenuation emerged more robustly in a simpler, two-target environment. Additional analyses revealed that reward accumulation predicted weaker learning and reduced the variability of corrective adjustments, suggesting that reward stabilizes motor output and downregulates sensitivity to SPEs. Collectively, our results demonstrate that implicit adaptation is not isolated from reward or motivational signals. Rather, reward appears to modify how the motor system interprets or weights SPEs, particularly when the task structure allows a consistent mapping between action and rewarded outcomes.

The primary motivation for this work was the observation that most previous studies examining how reward influences adaptation have operationalized reward in terms of successful target hits (Al-Fawakhiri et al., 2023; Kim et al., 2019; Oza et al., 2024). In these designs, “reward” is obtained only when the cursor lands on or close to the target, meaning that rewarded trials are also trials in which the SPE is small or absent. As a result, any reduction in learning following rewarded trials could simply reflect the limited error available to drive adaptation. Furthermore, successful performance may alter credit-assignment and state-estimation processes, reducing uncertainty about the executed motor command and changing how the system weights the observed error (Dam et al., 2013; Wei & Körding, 2009). These success-related changes, rather than reward itself, could therefore explain the observed modulation of implicit learning. The present experiments were designed to remove this ambiguity; we delivered explicit reward even when the cursor unambiguously missed the target. This dissociation between task outcome and reward (Vassiliadis et al., 2021) ensured that the effects we observed reflect a genuine impact of externally delivered reward. In this sense, the current findings extend past work by showing that reward can directly gate sensitivity to a fixed SPE, rather than indirectly shaping adaptation through success-related cues.

How might reward interact with mechanisms underlying implicit adaptation? One possibility is that reward might reduce the weight assigned to the SPE when updating motor commands (van der Kooij & Overvliet, 2016). Under Bayesian and Kalman filter based frameworks (Burge et al., 2008; Körding & Wolpert, 2004; Wei & Körding, 2009), reward could act as an additional form of evidence that the executed motor plan was appropriate, thereby decreasing the Kalman gain applied to error. Obtaining reward may thus signal that the SPE is less informative or less behaviorally relevant, leading to smaller updates. A second, complementary possibility is that reward decreases sensorimotor variability and promotes exploitation of the current policy (Pekny et al., 2015; Therrien et al., 2016; Uehara et al., 2019). The reduction in STL and inter-trial variability seen in the present work is consistent with this view. In this framework, reward may strengthen the current motor plan, suppressing deviations that would otherwise be expressed through implicit adaptation. Finally, it is possible that reward affects the expression of the acquired memory rather than the memory itself. If learning and its behavioral expression are indeed dissociable as some studies suggest (Heald et al., 2021; Hutter & Taylor, 2018; Schaefer et al., 2012; Taylor et al., 2014), then the underlying memory may continue to accumulate but may be expressed less strongly following rewarded trials. Although our current results cannot conclusively distinguish between these accounts, they converge on the idea that reward modulates implicit learning without altering the sensory input that generates the SPE.

Although reward consistently attenuated single-trial corrections in both experiments, a key difference emerged at the block level. In the 8-target experiment, reward did not significantly influence the overall magnitude or time course of adaptation, despite its reliable influence on STL. In contrast, in the 2-target experiment, these single-trial effects accumulated to produce robust group differences at asymptote and during washout. It is plausible that when target location varies from trial to trial as in Experiment 1, the small reward-induced reduction of STL (12-18%) is distributed across several movement directions, reducing its impact on any individual target. In contrast, when movements are made repeatedly to only two directions (Experiment 2), the same reward-modulated correction is applied to a stable motor plan, allowing small, trial-level differences to accumulate. This interpretation is consistent with studies showing that consistent task structure promotes both stronger error-based as well as reward-based adaptation (Darshan et al., 2014; Huberdeau et al., 2015; van der Kooij & Smeets, 2019). In this sense, the simplified 2-target environment provides the consistency needed for reward to exert a sustained influence on implicit learning.

Generalization characteristics of implicit adaptation could provide an additional explanation for the absence of group differences in Experiment 1. Implicit adaptation generalizes well to nearby movement directions (Donchin et al., 2003; Morehead et al., 2017; Taylor & Ivry, 2013). As a result, SPE-driven updates induced at one target inevitably influence movements to neighboring targets. In the 8-target task, such generalized updates can shift hand angles away from adjacent targets on subsequent trials, thereby reducing the likelihood that the unseen hand will successfully intersect those targets and receive reward. Consequently, although an SPE is present on every clamp trial, reward becomes sparse across directions. Indeed, participants in Experiment 1 received reward on only ∼24-32% of trials, whereas reward was obtained on ∼52-58% of trials in Experiment 2. With such sparse reinforcement, reward cannot consistently reinforce a target-specific motor plan in the 8-target task, limiting the accumulation of reward effects (Darshan et al., 2014). This explanation also aligns with findings from van der Kooij and Smeets (2019) who showed that reward-based learning is highly sensitive to spatial complexity, and with broader work on credit assignment in multi-context environments (Dam et al., 2013). When movements span multiple directions, the motor system may require many more repetitions per target than were available in Experiment 1. With inconsistent reward and limited repetitions, reward perhaps cannot reinforce a specific motor plan, thereby preventing accumulation of reward effects. In contrast, the more stable 2-target environment supports consistent reward-action associations, enabling small STL modulations to accumulate over time.

The present results open several avenues for future work. The contrast between the 8-target and 2-target designs suggests that task structure is critical in determining the influence of reward on SPE-driven learning. Future studies could systematically manipulate the spatial separation of targets to determine how it shapes the extent to which reward modulates error sensitivity. Similarly, using probabilistic reward schedules, or controlling reward frequency, may help isolate the contribution of reward properties from its binary presence or absence. Additionally, modeling frameworks that jointly estimate reward-based and error-based learning rates (Cashaback et al., 2017; Izawa & Shadmehr, 2011; Thrommershäuser et al., 2008) may help formalize how these signals are combined. Studies that include cerebellar or parietal stimulation, or studies of patients with dopaminergic dysfunction (e.g., Parkinson’s disease), may help clarify whether the interaction between reward and adaptation arises within systems that drive adaptation or reinforcement learning circuits, or their interaction. Together, these studies may help build a unified account of how reward and error jointly shape motor learning.

## Acknowledgements

This work was supported by grants from the Department of Science and Technology, Government of India. Support from IIT Gandhinagar is also acknowledged.

